# Proximal relationships of moonlighting Proteins in *Escherichia coli*: a mathematical genomic perspective

**DOI:** 10.1101/2025.01.16.633489

**Authors:** Debaleena Nawn, Sk. Sarif Hassan, Moumita Sil, Ankita Ghosh, Arunava Goswami, Vladimir N. Uversky

## Abstract

Moonlighting proteins in *Escherichia coli* (*E.coli*) perform multiple independent functions without altering their primary amino acid sequence, challenging the “one gene-one enzyme” hypothesis. Bacterial proteins serve various functions, including host cell adhesion, extracellular matrix interaction, and immune modulation, while also supporting essential physiological processes within the bacteria. Identifying these proteins in pathogens and tracking their genetic changes is crucial for understanding bacterial survival and virulence. A quantitative understanding of these proteins is pivotal as it enables the identification of specific patterns and relationships between amino acid composition, protein stability, and functional versatility. This study quantitatively analyzes fifty *E. coli* moonlighting proteins, focusing on their structural and functional features. Key findings include variability in amino acid composition, with alanine predominating, and a preference for non-polar residues, which may enhance protein stability. Quantitative features analyses identified seven distinct proximal sets, reflecting the proteins’ spatial arrangements of amino acids, structural diversity, and functional roles in processes such as metabolism, stress response, and gene regulation. These results deepen our understanding of the multifunctionality of *E. coli* moonlighting proteins, indicating their adaptability and implications for bacterial survival and pathogenicity.

## 1. Introduction

Moonlighting proteins are proteins that voluntarily juggle other distinct functions alongside their primary role without undergoing any changes in their primary amino acid sequence. The concept of moonlighting in proteins, originally termed gene sharing by Piatigorsky, has evolved into the widely accepted term “moonlighting”, akin to individuals who perform multiple jobs [1, 2]. These proteins are unique in the realm of multi-functionality, as they carry out several independent and often unrelated functions without compartmentalizing these activities into separate domains within the protein [3]. The term “moonlighting proteins” specifically excludes certain categories of proteins [4]. It does not apply to homologous proteins that are not identical, nor to proteins that exhibit multiple functions due to gene fusions, splice variants, or different proteolytic fragments. Additionally, proteins that carry out the same function in various cellular locations are not considered moonlighting proteins. This distinction emphasizes that moonlighting proteins uniquely perform multiple, independent functions without these forms of genetic or structural variations [2, 5]. Also, it is important to differentiate moonlighting proteins from pleiotropic effects. Pleiotropy occurs when a single protein’s function impacts various cellular processes, such as a protein interacting with multiple partners in different pathways or an enzyme playing a role in several metabolic pathways [6]. In contrast, moonlighting proteins carry out distinct functions that are mechanistically different from one another [7].

In the context of protein moonlighting, the hypothesis proposed by Beadle and Tatum—originally stated as “one gene-one enzyme” and later refined to “one gene-one polypeptide”—is significantly challenged [8]. Beadle and Tatum’s hypothesis suggested a direct and singular relationship between a gene and its corresponding protein product, which performs a specific function [9]. However, with the discovery of protein moonlighting, it is clear that a single protein can have multiple roles, thus complicating the one gene-one polypeptide-one function paradigm. This modern understanding illustrates that proteins are not restricted to a single function. Instead, they can participate in various biochemical pathways and processes, reflecting a higher level of complexity in genetic and proteomic interactions than originally proposed by Beadle and Tatum [9]. This complexity is particularly evident in more evolved organisms, where proteins often exhibit multi-functionality to support intricate physiological processes without a proportional increase in genome size [4, 10].

Previous studies have identified over 800 moonlighting proteins, many of which exhibit conserved additional functions [10, 11, 12, 13]. Notably, glycolytic enzymes are multifunctional, while enzymes in other pathways typically have only one additional function [14]. These proteins are involved in various cellular processes, such as apoptosis, gene regulation, transport, and signaling, which contribute to cellular energy efficiency [15]. In prokaryotes, their adaptability aids in evading host immune responses [7, 16]. Moonlighting proteins are particularly widespread in bacteria, providing a unique advantage by enabling multiple functions from the same genetic material, making them promising targets for combating drug resistance [17]. Additionally, these proteins act as bacterial adhesins, facilitating interactions with the environment and responses to environmental changes by binding to host cells, the cytoskeleton, and immune system components [18].

Despite the indication that many organisms’ genomes contain a vast array of these proteins playing crucial roles in various pathways and disease processes, the current list of verified moonlighting proteins remains too limited to fully grasp the complex cellular mechanisms underlying their diverse functions [19, 20, 21, 22]. Our previous study took a quantitative informatics approach to analyze human moonlighting protein sequences, providing insights into their evolutionary connections [22]. Phylogenetic analyses of these proteins uncover distal-proximal relationships through sequence-derived quantitative features [22]. Previous proteomics studies have highlighted challenges in identifying the functions of moonlighting proteins, as sequence homology does not always correlate with functional attributes [23]. While proteomics can identify moonlighting proteins with significant structural or sequence differences, it struggles with those that share structural homology. In such cases, automated genome annotation methods may fail to accurately determine their functions [24]. Therefore, the analytical approach used in this study, along with previous studies on human moonlighting proteins, is crucial for defining and categorizing these proteins into distinct functional groups [22]. This study explores the variation in the world of moonlighting proteins in *Escherichia coli*. The moonlighting functions of bacterial proteins are associated with, but not restricted to, processes such as adhesion to host epithelial cells, interaction with the host extracellular matrix, and modulation of the host immune response [25, 26]. Identifying these proteins in pathogenic bacteria and tracking their genetic changes over time is crucial for understanding their role in bacterial survival and virulence, particularly in the context of multidrug resistance [27].

Despite the growing recognition of moonlighting proteins and their roles in various pathways and diseases, the current list of verified proteins remains limited, hindering a comprehensive understanding of their multi-functionalities [19, 20, 21, 22]. Our previous study utilized a quantitative informatics approach to analyze human moonlighting protein sequences, revealing distal-proximal relationships [22]. Proteomics studies have highlighted the challenges in identifying these proteins, as sequence homology does not always correlate with functional attributes [23]. While proteomics can detect proteins with significant structural differences, it struggles with those that share structural homology, and automated genome annotation methods may fail to accurately determine their functions [24]. Thus, the analytical approach applied in this study is essential for defining and categorizing these proteins into distinct functional groups [22]. This research also explores moonlighting proteins in Escherichia coli, focusing on processes like adhesion to host epithelial cells, interaction with the extracellular matrix, and modulation of the host immune response [25, 26]. Tracking these proteins and their genetic changes is crucial for understanding their role in bacterial survival and virulence, particularly in the context of multidrug resistance [27].

The quantitative shortfall in identifying moonlighting proteins is largely due to the serendipitous discovery of their additional functions in unrelated experimental contexts [24]. The importance of quantitative genomics in the study of *Escherichia coli* moonlighting proteins lies in its ability to provide a comprehensive view of how these proteins evolve and diversify within the genome, offering insights into their functional and regulatory networks. This study explores quantitative proximal relationships among *Escherichia coli* moonlighting proteins, focusing on the connection between sequence-based homology and their multifaceted functions.

## 2. Data

In the present study, fifty promiscuous proteins/moonlighting proteins of *Escherichia coli* (*E.coli*) were extracted from the database MultitaskProtDBII (Table 1) [28].

**Table 1:**
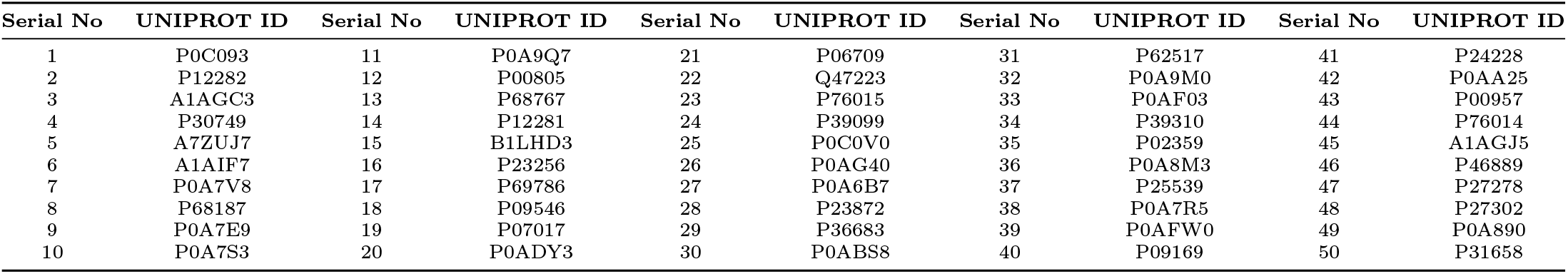
List of Moonlighting proteins in *E.coli* with their associated Uniprot ID (Hyperlinked with respective Uniprot webpage)

**Table 2:**
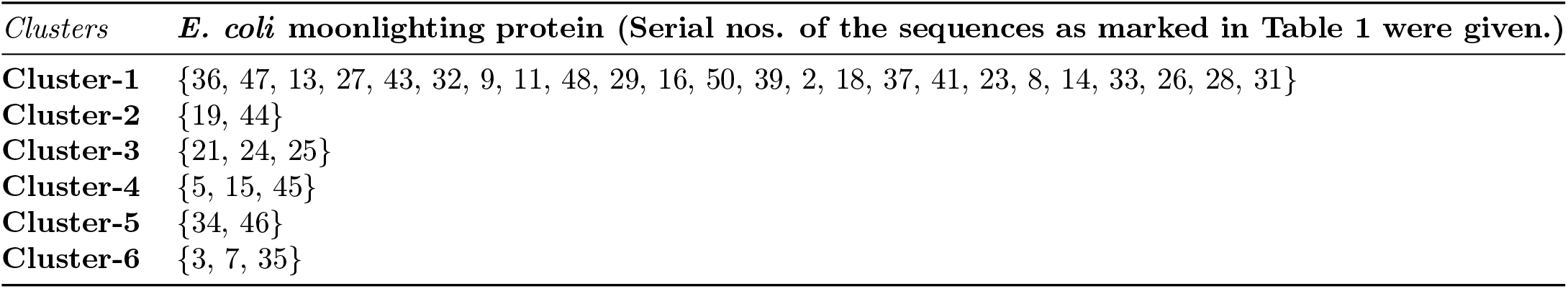
Clusters derived from relative frequency of amino acids.

| Clusters | *E. coli* moonlighting protein (Serial nos. of the sequences as marked in Table 1 were given.) |
| --- | --- |
| **Cluster-1** | {36, 47, 13, 27, 43, 32, 9, 11, 48, 29, 16, 50, 39, 2, 18, 37, 41, 23, 8, 14, 33, 26, 28, 31} |
| **Cluster-2** | {19, 44} |
| **Cluster-3** | {21, 24, 25} |
| **Cluster-4** | {5, 15, 45} |
| **Cluster-5** | {34, 46} |
| **Cluster-6** | {3, 7, 35} |

**Table 3:** Frequency of homogeneous poly-string of length 1, 2, … 6 of *E.coli* moonlighting proteins.

| Serial No. | Uniprot ID | Homogenous polystring of length l |  |  |  |  | Serial No. | Uniprot ID | Homogenous polystring of length l |  |  |  |  |
| --- | --- | --- | --- | --- | --- | --- | --- | --- | --- | --- | --- | --- | --- |
|  |  | l=1 | l=2 | l=3 | l=4 | l=6 |  |  | l=1 | l=2 | l=3 | l=4 | l=6 |
| 1 | P0C093 | 172 | 11 | 0 | 1 | 0 | 26 | P0AG40 | 276 | 17 | 1 | 0 | 0 |
| 2 | P12282 | 231 | 9 | 0 | 0 | 0 | 27 | P0A6B7 | 358 | 20 | 2 | 0 | 0 |
| 3 | A1AGC3 | 112 | 6 | 2 | 0 | 0 | 28 | P23872 | 266 | 25 | 1 | 0 | 0 |
| 4 | P30749 | 138 | 6 | 0 | 0 | 0 | 29 | P36683 | 744 | 56 | 3 | 0 | 0 |
| 5 | A7ZUJ7 | 210 | 12 | 0 | 0 | 0 | 30 | P0ABS8 | 73 | 0 | 1 | 0 | 0 |
| 6 | A1AIF7 | 145 | 7 | 2 | 0 | 0 | 31 | P62517 | 746 | 43 | 3 | 0 | 1 |
| 7 | P0A7V8 | 184 | 11 | 0 | 0 | 0 | 32 | P0A9M0 | 700 | 39 | 2 | 0 | 0 |
| 8 | P68187 | 336 | 16 | 1 | 0 | 0 | 33 | P0AF03 | 170 | 11 | 1 | 0 | 0 |
| 9 | P0A7E9 | 212 | 13 | 1 | 0 | 0 | 34 | P39310 | 181 | 14 | 1 | 0 | 0 |
| 10 | P0A7S3 | 115 | 3 | 1 | 0 | 0 | 35 | P02359 | 158 | 9 | 1 | 0 | 0 |
| 11 | P0A9Q7 | 784 | 44 | 5 | 1 | 0 | 36 | P0A8M3 | 583 | 28 | 1 | 0 | 0 |
| 12 | P00805 | 303 | 21 | 1 | 0 | 0 | 37 | P25539 | 325 | 21 | 0 | 0 | 0 |
| 13 | P68767 | 449 | 24 | 2 | 0 | 0 | 38 | P0A7R5 | 101 | 1 | 0 | 0 | 0 |
| 14 | P12281 | 373 | 16 | 2 | 0 | 0 | 39 | P0AFW0 | 152 | 2 | 2 | 0 | 0 |
| 15 | B1LHD3 | 174 | 12 | 1 | 0 | 0 | 40 | P09169 | 268 | 23 | 1 | 0 | 0 |
| 16 | P23256 | 357 | 15 | 1 | 0 | 0 | 41 | P24228 | 435 | 18 | 2 | 0 | 0 |
| 17 | P69786 | 409 | 31 | 2 | 0 | 0 | 42 | P0AA25 | 100 | 3 | 1 | 0 | 0 |
| 18 | P09546 | 1167 | 69 | 5 | 0 | 0 | 43 | P00957 | 763 | 46 | 7 | 0 | 0 |
| 19 | P07017 | 459 | 41 | 2 | 0 | 1 | 44 | P76014 | 182 | 14 | 0 | 0 | 0 |
| 20 | P0ADY3 | 99 | 9 | 2 | 0 | 0 | 45 | A1AGJ5 | 112 | 9 | 0 | 0 | 0 |
| 21 | P06709 | 283 | 16 | 2 | 0 | 0 | 46 | P46889 | 1057 | 117 | 10 | 2 | 0 |
| 22 | Q47223 | 150 | 17 | 0 | 0 | 0 | 47 | P27278 | 353 | 27 | 1 | 0 | 0 |
| 23 | P76015 | 303 | 22 | 3 | 0 | 0 | 48 | P27302 | 600 | 30 | 1 | 0 | 0 |
| 24 | P39099 | 397 | 26 | 2 | 0 | 0 | 49 | P0A890 | 67 | 7 | 0 | 0 | 0 |
| 25 | P0C0V0 | 394 | 34 | 4 | 0 | 0 | 50 | P31658 | 242 | 19 | 1 | 0 | 0 |

## 3. Methods

### 3.1. Determining amino acid frequency composition

Amino acid frequency, defined as the count of each amino acid in a protein sequence, was computed for all 50 *E. coli* moonlighting proteins [29, 30, 31]. To account for the varying lengths of the moonlighting protein sequences, the relative frequency of each amino acid was calculated by dividing the amino acid frequency by the length of the respective sequence and multiplying by 100. This relative frequency represents the proportion of each amino acid in the sequence. For each protein sequence, the relative frequencies of the 20 amino acids form a 20-dimensional vector.

### 3.2. Determining homogeneous poly-string frequency of amino acids

A homogeneous poly-string of length *n* is defined as a sequence of *n* consecutive occurrences of a particular amino acid [32]. For example, the sequence “SSSWWSWW” contains one homogeneous poly-string of S with length 3, one homogeneous poly-string of S with length 2, and two homogeneous poly-strings of W with length 2. When counting homogeneous poly-strings of a given length *n*, only the exclusive occurrences of length *n* are considered. To evaluate these poly-strings across all protein sequences, we first computed the maximum possible length of homogeneous poly-strings among all amino acids in the dataset. Based on this maximum length, we then enumerated the counts of homogeneous poly-strings for each amino acid in each sequence, considering all possible lengths from 1 up to the maximum length observed.

### 3.3. Evaluating polar, non-polar residue profiles

Each amino acid in a given *E. coli* moonlighting protein sequence was classified as either polar (P) or non-polar (Q). This classification converted the protein sequence into a binary sequence consisting of two symbols: P and Q. The resulting binary P-Q profile provides a representation of the spatial arrangement of polar and non-polar residues across the protein sequence, allowing for the analysis of their distribution and periodicity [33, 34, 35].

#### 3.3.1. Change response sequences based on polar, non-polar residue profiles

In the binary polar-non-polar profile of each moonlighting protein sequence, four possible changes can occur between two consecutive residues: Polar to Polar (PP), Polar to Non-polar (PN), Non-polar to Non-polar (NN), and Non-polar to Polar (NP). These changes were recorded as a sequence that reflects the spatial arrangement of polar and non-polar residues in the P-Q binary profile. This sequence is referred to as the “P-Q Change Response Sequence” (*CRS*_*P Q*_). The frequency of each of the four transition types (PP, PN, NN, and NP) was then calculated for each moonlighting protein, based on its corresponding binary polar-non-polar profile.

### 3.4. Evaluating acidic, basic, neutral residue profiles

Each amino acid in a given moonlighting protein sequence was classified as acidic (A), basic (B), or neutral (N). This classification transformed the protein sequence into a ternary sequence, referred to as the A-B-N profile, consisting of three symbols: A, B, and N. This ternary profile captures the spatial distribution of acidic, basic, and neutral residues across the protein sequence, providing insights into its overall charge and polarity characteristics.

#### 3.4.1. Change response sequences based on acidic-basic-neutral residue profiles

In the A-B-N ternary profile of each moonlighting protein sequence, there are nine possible changes between two consecutive residues: Acidic to Acidic (AA), Acidic to Basic (AB), Acidic to Neutral (AN), Basic to Acidic (BA), Basic to Basic (BB), Basic to Neutral (BN), Neutral to Acidic (NA), Neutral to Basic (NB), and Neutral to Neutral (NN). These transitions were recorded in sequence according to the spatial arrangement of acidic, basic, and neutral residues within the A-B-N profile. This sequence is referred to as the “A-B-N Change Response Sequence (*CRS*_*ABN*_)”. The frequency of each of the nine possible transitions (AA, AB, AN, BA, BB, BN, NA, NB, and NN) was then counted for each moonlighting protein, based on its corresponding A-B-N ternary profile.

### 3.5. Evaluating intrinsic protein disorder

The predisposition for intrinsic disorder in all human moonlighting proteins analyzed in this study was assessed using a set of well-established per-residue disorder predictors, including PONDR® VLS2, PONDR® VL3, PONDR® VLXT, PONDR® FIT, IUPred-Long, and IUPred-Short [36, 37, 38, 39, 40, 41]. The Rapid Intrinsic Disorder Analysis Online (RIDAO) platform was utilized to gather the results from each predictor in bulk [42].

Per-residue disorder scores range from 0 (fully ordered) to 1 (fully disordered), with residues scoring above 0.5 considered disordered (D). Residues with scores between 0.25 and 0.5 were classified as highly flexible (HF), while those between 0.1 and 0.25 were classified as moderately flexible (MF) [41]. Residues with scores less than 0.1 were termed as ‘other’ (O).

#### 3.5.1. Change response sequences based on intrinsic protein disorder residues

There are sixteen possible transitions between two consecutive residues in the sequence, including: (HF_D, HF_HF HF_MF, HF_O, MF_D, MF_HF, MF_MF, MF_O, O_D, O_HF, O_MF, and O_O). These transitions were recorded in sequence according to the spatial arrangement of D, HF, MF, and O residues in each moonlighting protein sequence. The frequency of each of the sixteen possible transitions was then counted from the change response sequences generated for each protein, providing insight into the spatial dynamics of protein flexibility and disorder across the sequences.

### 3.6. Formation of distance matrices and dendrograms

Euclidean distance was computed between the feature vectors of all pairs of moonlighting protein sequences for each of the following four features:

- Relative frequency of amino acids (dimension 20)
- Relative frequency of changes from polar-nonpolar profiles (dimension 4)
- Relative frequency of changes from acidic-basic-neutral profiles (dimension 9)
- Relative frequency of changes from disordered, highly flexible, moderately flexible, and other residues (dimension 16)

Relative frequency was defined as the frequency of each feature divided by the length of the sequence and multiplied by 100 [22]. Dendrograms were constructed based on the Euclidean distances using average linkage. Different color thresholds (empirically chosen) were applied to the dendrograms, where a threshold value *T* determined the color assignment for each group of nodes. Any group of nodes with a linkage value less than *T* was assigned a unique color in the dendrogram.

### 3.7. Proximal relationships of moonlighting proteins

*E. coli* moonlighting protein sequences belong to the same cluster in all dendrograms are termed as *proximal* sequences. To identify the final proximal sequences, we focused on proteins that were clustered together in all four features. The intersection of these clusters was calculated, and only those moonlighting proteins that were consistently clustered together across all features were selected.

## 4 Results and analyses

### 4.1. Relative frequency of amino acids in E.coli moonlighting protein sequences

From the relative frequency vector (consisting of 50 elements) of each amino acid, maximum and minimum median was recorded. As the amino acid distribution was skewed, the median value was considered instead of average (Figure1). Alanine had the highest median (10.1) followed by Leucine (9.2) and Cysteine (0.79) had the lowest median followed by Tryptophan (0.87). Maximum standard deviation was found for alanine (2.85) followed by Arginine (2.79) while minimum was noted for Cysteine (0.86). Based on the percentage of amino acids in individual sequences, the maximum value was observed for alanine in the sequence A1AIF7 (20) followed by Q47223 (17.39). Tryptophan, Cysteine, and Histidine was absent in 13, 11, and 1 number of sequences, respectively.

**Figure 1:**
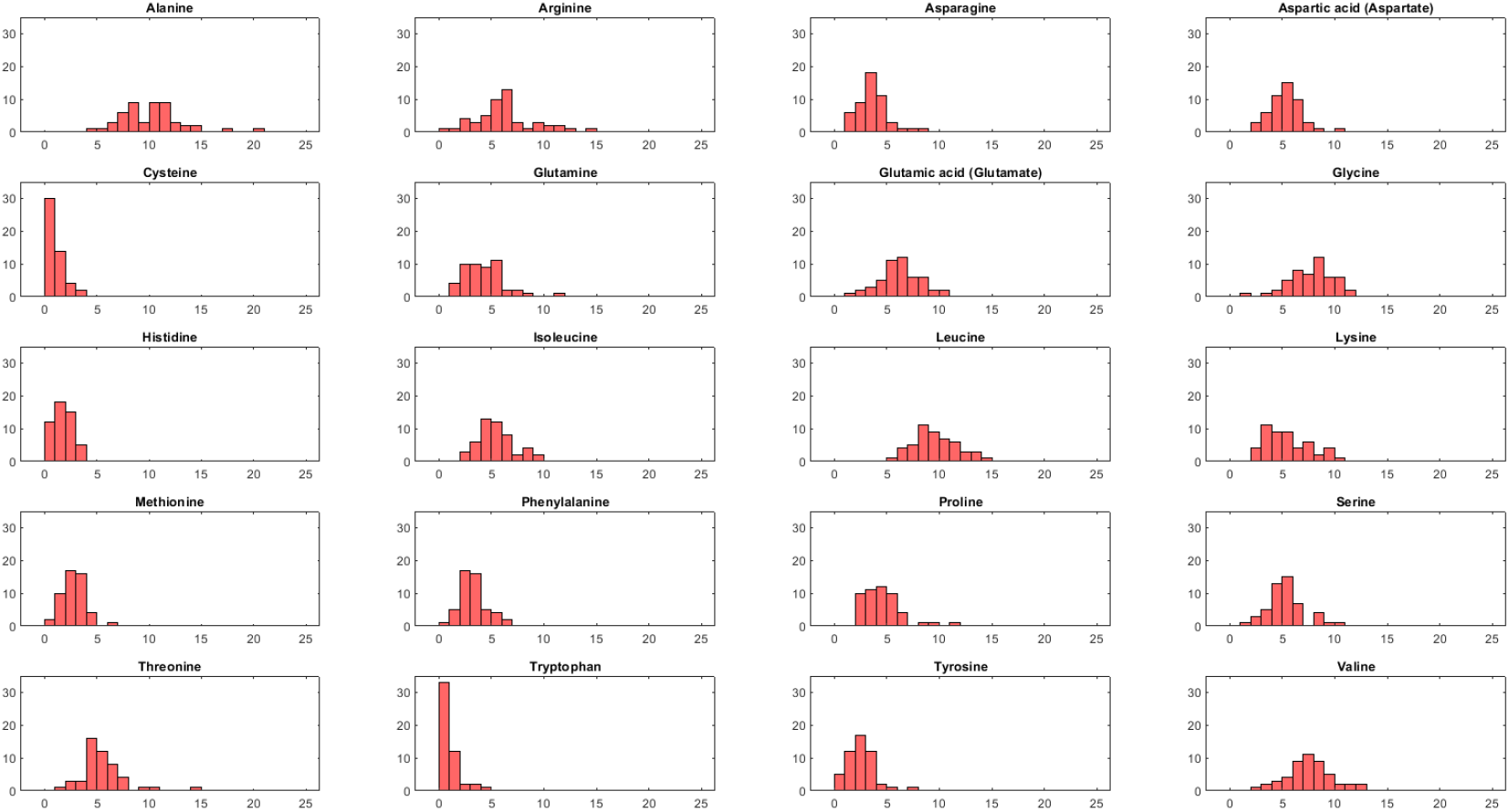
Histogram of relative frequency of each amino acid. X axis denotes percentage and Y axis denotes number of sequences

Based on a threshold of distance of 9, 37 moonlighting sequences formed 6 clusters as obtained in the dendrogram (Figure 2). The largest cluster comprised of 24 sequences as colored in green (Figure 2 and Table 5). Furthermore, it was observed that the sequence Q47223 (Serial No. 22) was an outlier, as depicted from the dendrogram.

**Figure 2:**
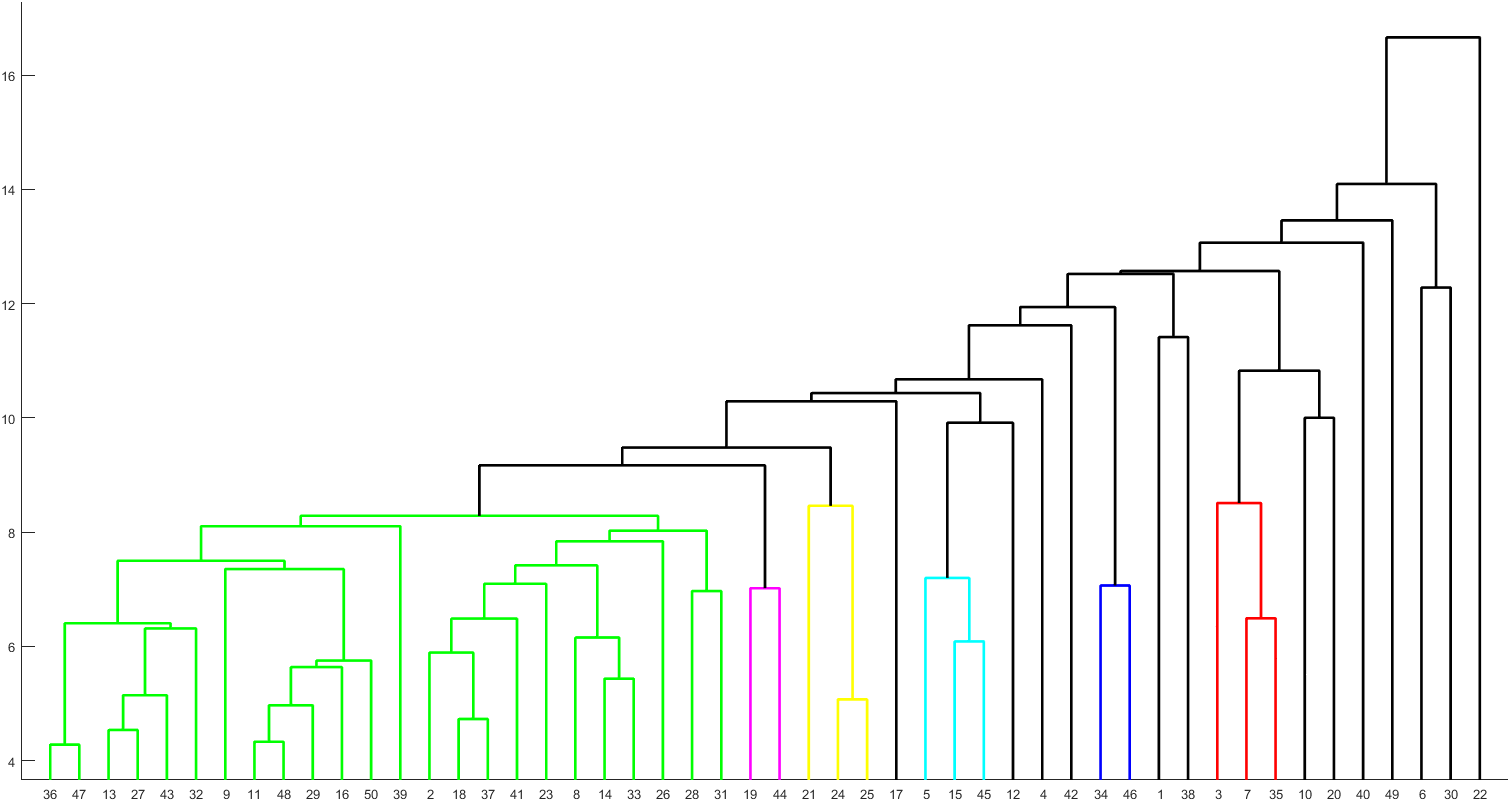
Phylogenetic relationship among the moonlighting proteins based on relative frequency of amino acids.

### 4.2. Homogeneous poly-string frequency of amino acids in E.coli moonlighting protein sequences

The maximum length of a homogeneous poly-string found to be 6, considering all amino acids across 50 sequences. Hence, frequency of homogeneous poly-strings with lengths ranging from 1 to 6 for each of the twenty amino acids in each sequence were tabulated (**Supplementary file-1**). The total count of poly-strings across all amino acids in each sequence was summarized in Table 4. No poly-string of length 5 was found in any sequence (Refer to **Supplementary file-1**).

**Table 4:** Clusters of moonlighting proteins based on relative frequency of PP, NP, PP, and NN changes as obtained from Polar, non-polar profiles.

| Clusters | Moonlighting sequences |
| --- | --- |
| Cluster 1 | {36, 47, 19, 35, 30, 3, 10, 12, 43, 34, 15, 32} |
| Cluster 2 | {7, 38} |
| Cluster 3 | {2, 9, 6, 24, 14, 26, 29, 42, 8, 16, 33, 37, 18, 23, 45, 48, 25, 21, 31} |
| Cluster 4 | {4, 11, 28, 20, 13, 44, 41, 46, 50, 49} |
| Cluster 5 | {5, 22, 27, 39} |

**Table 5:** Clusters of moonlighting proteins based on relative frequency of BA, NA, AA, BB, NB, AB, BN, NN, and AN changes as obtained from acidic, basic, and neutral profiles.

| Clusters | Sequences |
| --- | --- |
| Cluster-1 | {12, 24, 31, 2, 14, 19, 34, 16, 18, 50, 44, 28, 46, 33, 23, 8, 11, 13, 39, 21, 48, 9, 29, 37, 40} |
| Cluster-2 | {25, 41} |
| Cluster-3 | {1, 6, 15, 45, 20} |
| Cluster-4 | {3, 35} |
| Cluster-5 | {30, 38} |
| Cluster-6 | {4, 36, 32, 27, 47, 42, 43} |
| Cluster-7 | {5, 26} |
| Cluster-8 | {17, 22} |

Poly-strings length of 6 were observed in *P* 07017 and *P* 62517, composed of alanine and arginine respectively, each with a frequency of 1. Poly-strings of length 4 appeared in sequences *P* 0*C*093, *P* 0*A*9*Q*7, and *P* 46889, consisting of leucine, valine and alanine, respectively, where former two had frequency of 1 and the last one had frequency of 2. 11 sequences did not show any poly-string of length 3. *P* 46889 had the highest number (10) of poly-string of length 3. None of the sequences contain poly-string of length 3 of following amino acids-Cysteine, Phenylalanine, Histidine, Tryptophan, and Tyrosine. 49 moonlighting proteins with homogeneous poly strings of length 2 was observed, only sequence *P* 0*ABS*8 did not have any poly string of length 2. None of the sequences contain WW (poly-string of Tryptophan having length 2).

### 4.3. Polar, non-polar residue profiles of E. coli moonlighting proteins

Figure 3 shows percentage distributions of polar and non-polar residues in each sequence. The average ratio of polar to non-polar residues was 0.89 *±* 0.14. P69786 (P09169) exhibited the lowest (the highest) ratio of 0.5 (1.28). Ratio was greater than 0.7 for all sequences except P69786. In P0ABS8, the ratio was 1 as the percentages of polar and non-polar residues were exactly identical.

**Figure 3:**
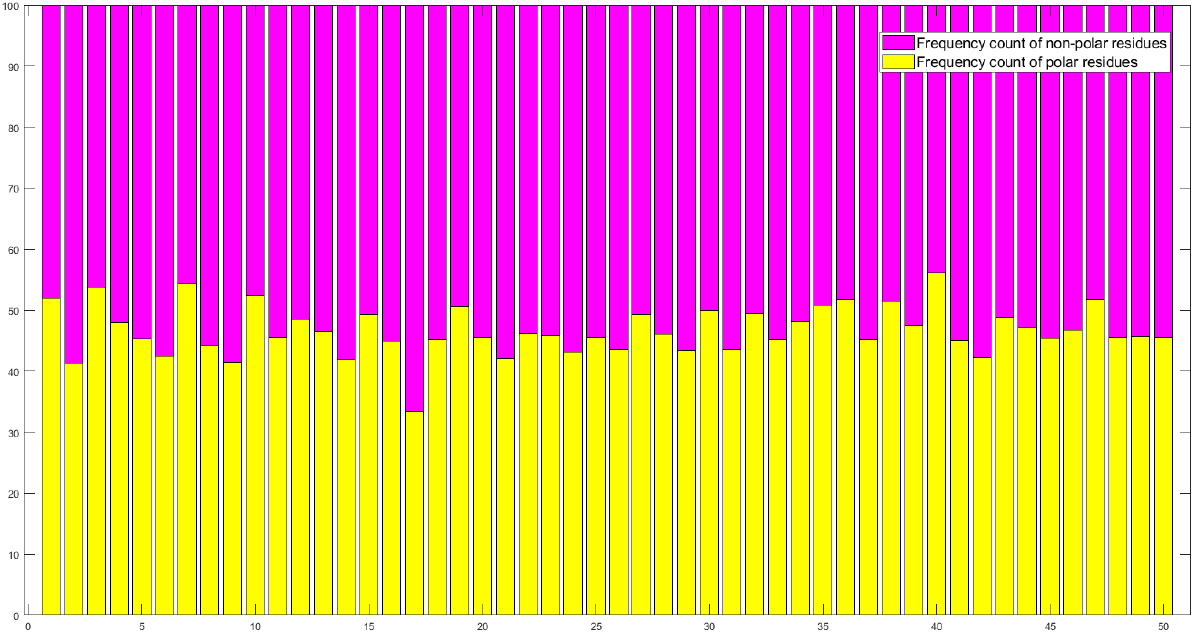
Percentages of polar, non-polar residues in *E. coli* moonlighting proteins

#### 4.3.1. Change response sequences of polar, non-polar profiles in E. coli moonlighting protein sequences

P69786 had the lowest percentages of ‘PN’ (19.95%), ‘NP’ (20.16%), and ‘PP’ (13.23%) changes (Figure 4). P09169 had the highest percentage of PP (32.28%) as it had the highest ratio of polar to non-polar residues. Percentages of NN changes for the sequences P69786 was significantly high (46.63%) compared to others as the ratio of non-polar to polar residue was 2 for this sequence.

**Figure 4:**
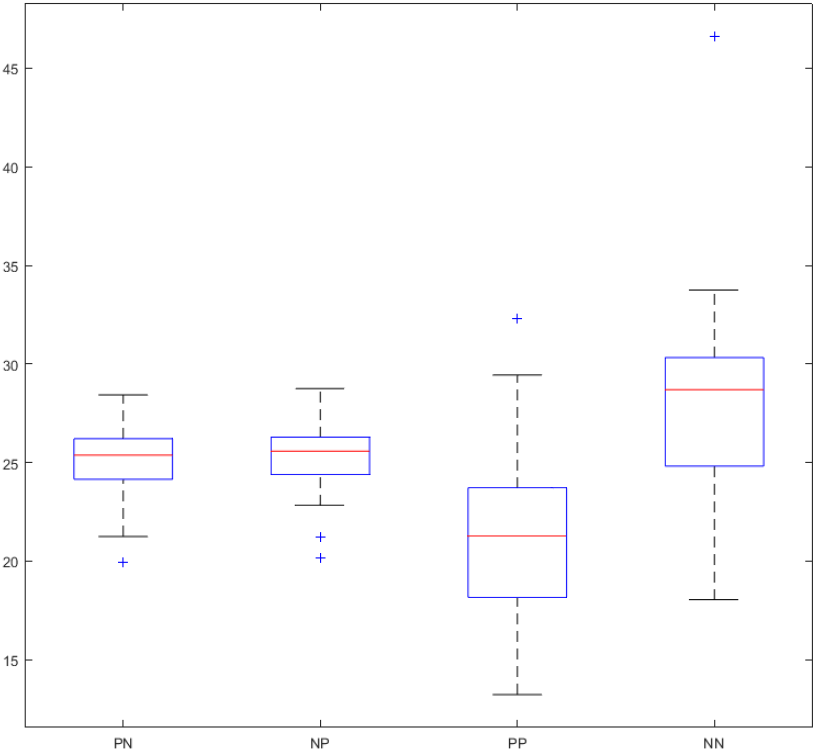
Box-plot of the relative frequency of PN, NP, PP, and NN changes in *E. coli* moonlighting proteins

Five clusters were formed based on a distance threshold of 5.6 (Figure 5 and Table 5). The largest cluster comprised of 19 sequences (Cluster 3), highlighted in light green in Figure 5. P69786 stood as outlier due to its distinct polar non-polar ratio.

**Figure 5:**
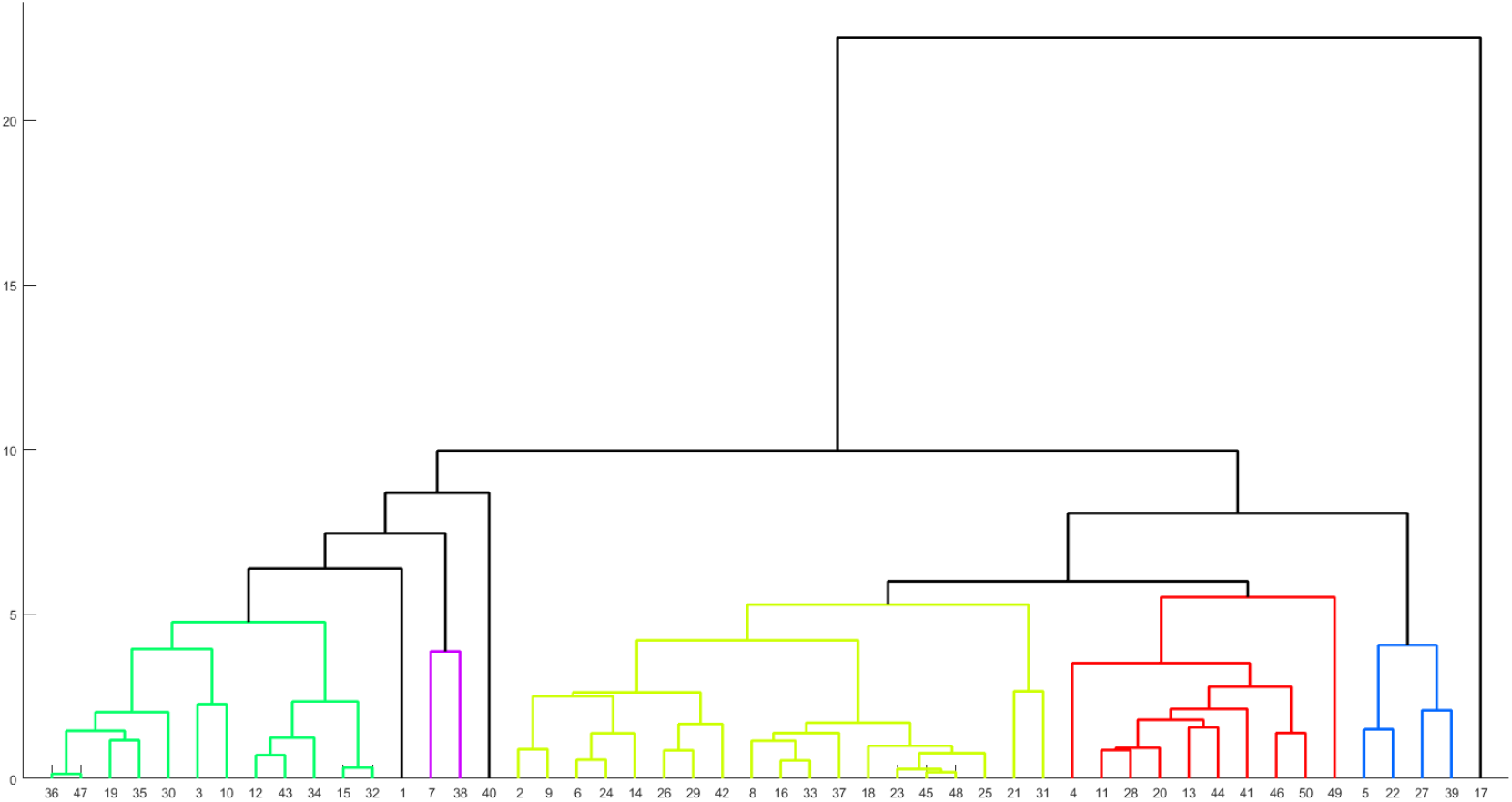
Phylogenetic relationship among the moonlighting proteins based on relative frequency of PP, NP, PP, and NN changes as obtained from polar, non-polar profiles.

### 4.4. Acidic, Basic, Neutral residue based phylogenetic relationship

Figure 6 exhibited percentages of acidic, basic and neutral residues in each sequence. Acidic, basic and neutral residues spanned from 5.64% (P0A7S3) to 16% (P30749), 5.43% (Q47223) to 22.58% (P0A7S3) and 67.47% (P0A7V8) to 88.04% (Q47223), respectively. Note that, associated sequences are mentioned in the parentheses.

**Figure 6:**
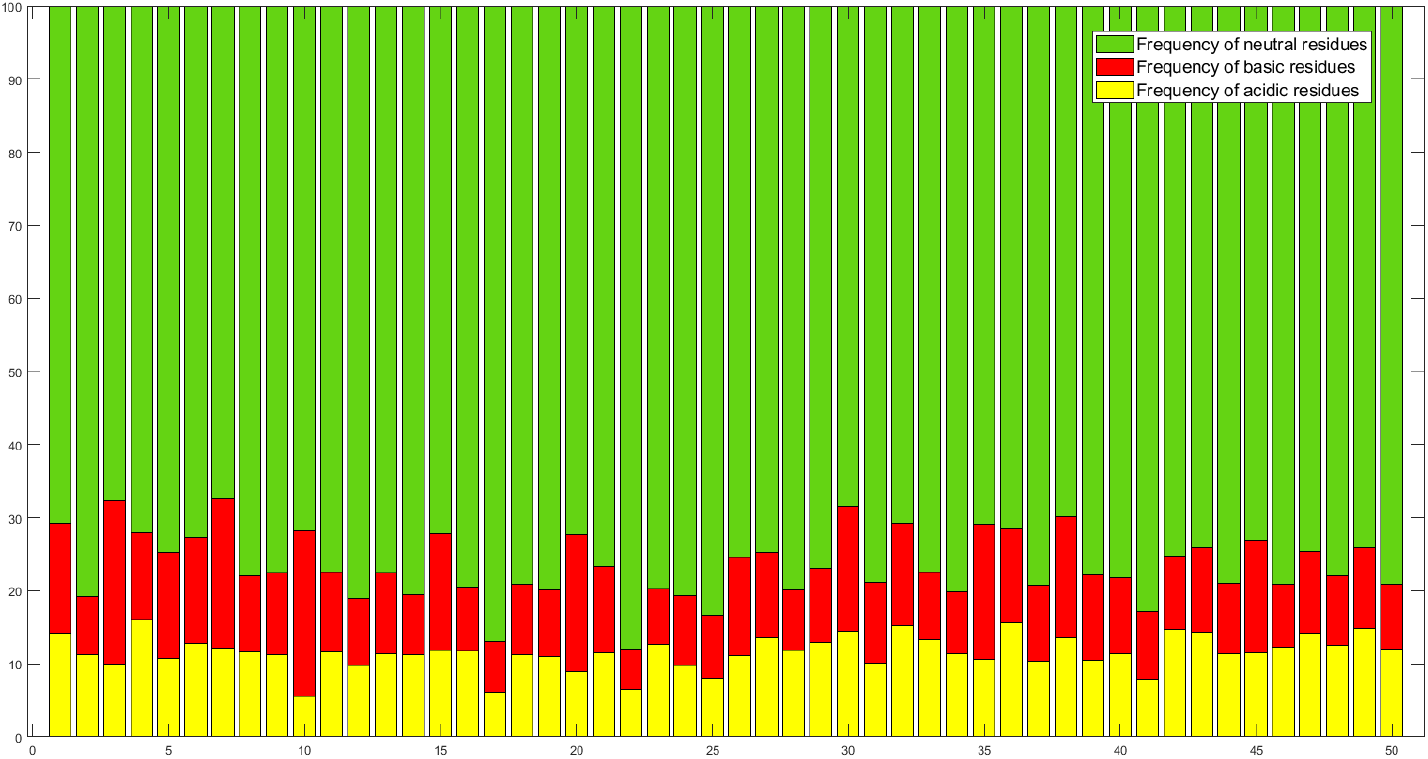
Percentages of acidic, basic, neutral residues in moonlighting proteins

#### 4.4.1. Change response sequences of acidic, basic, and neutral profiles

The relative frequency distribution of nine different residue changes, denoted as BA, NA, AA, BB, NB, AB, BN, NN, and AN, was visualized in Figure 7. NN changes varied from 42.92% (P30749) to 76.5% (Q47223). P30749 and A1AGC3 displayed the highest percentage of AA and BB changes respectively (3.35% and 6.98%). P0A7S3 stood out with the highest percentages of NB and BN changes (16.26% and 15.44% respectively), whereas P0ABS8 had the highest percentages of BA and AB(4% and 5.33%).

**Figure 7:**
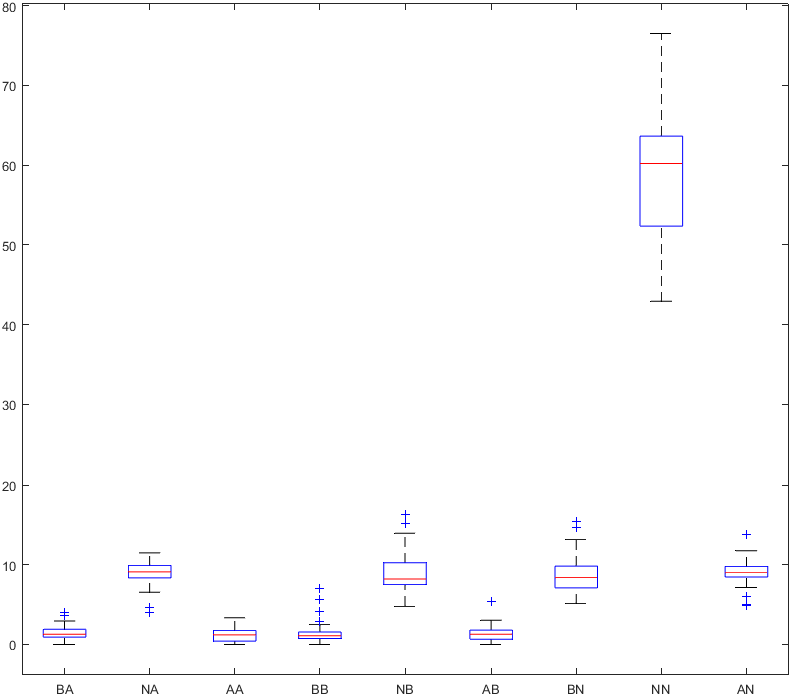
Box-plot of the relative frequency of all nine changes in moonlighting proteins

Following change responses were absent in following moonlighting sequences:

- Basic to Acidic: Q47223
- Acidic to Acidic: P0A7V8, P0A7E9, P0A7S3, P0ADY3, Q47223, P0ABS8, P0A7R5, P0AFW0, and P62328
- Basic to Basic: P12282, P23256, Q47223, P0ABS8, and P0AA25
- Acidic to Basic: P12282 and P0A890

Under a distance threshold of 5.1, the analysis led to the formation of eight distinct clusters comprising 48 moonlighting sequences. This information is visually depicted in the dendrogram (Figure 8) and summarized in Table 6. Notably, the largest cluster (Cluster 1) contained 25 sequences, depicted in light green colour in Figure 8. Specifically cluster consisting of sequences P06709 (17) and P0A7E9 (22) demonstrated considerable distance from others, reflecting their distinct positions within the dataset.

**Figure 8:**
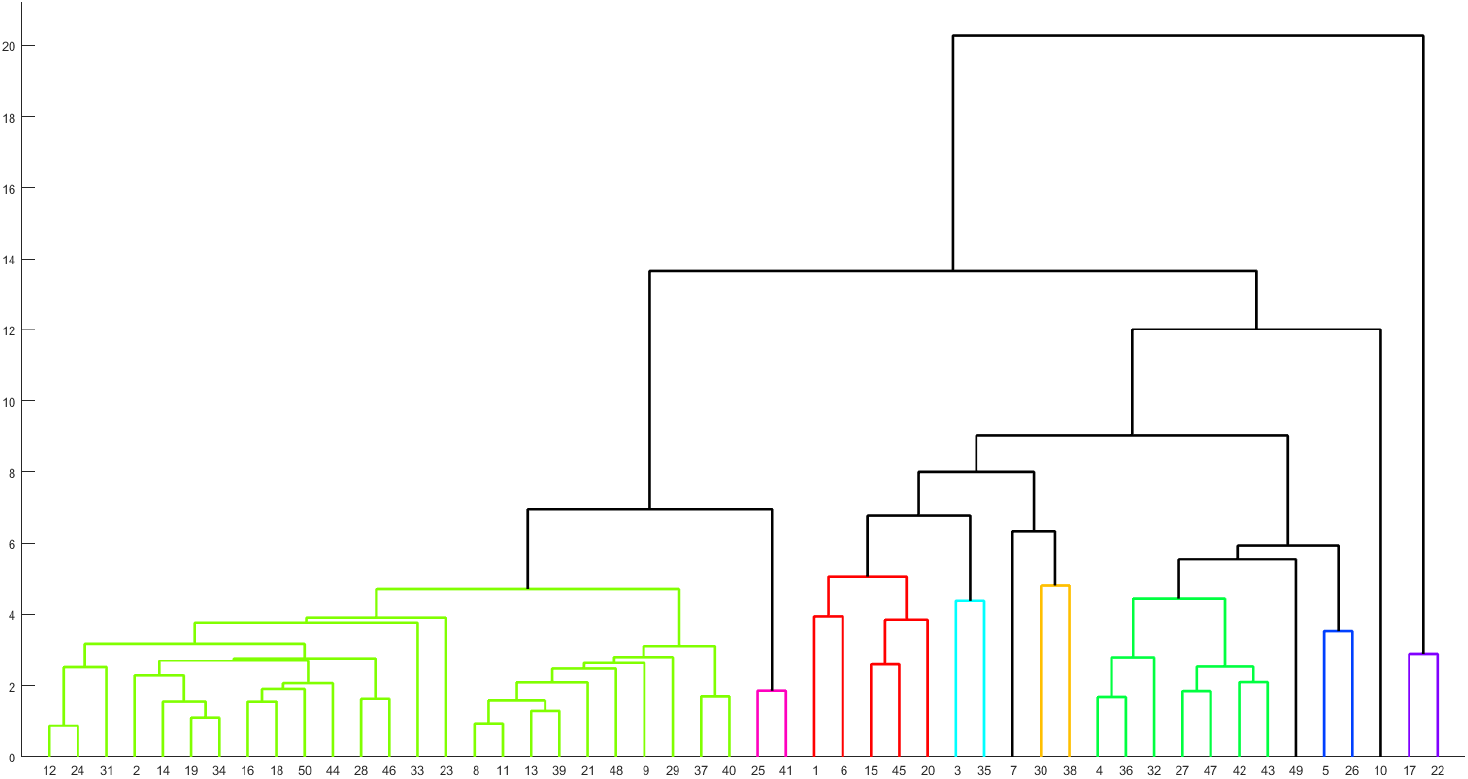
Phylogenetic relationship among the moonlighting proteins based on relative frequency of BA, NA, AA, BB, NB, AB, BN, NN, and AN changes as obtained from acidic, basic, and neutral profiles.

### 4.5. Intrinsic protein disorder analysis of E. coli moonlighting proteins

The percentages of disordered residues, highly flexible residues, moderately flexible residues, and other residues in each moonlighting protein were determined from the intrinsic protein disorder probability of each residue (Figure 9). Four moonlighting proteins exhibited a notably high percentage of disordered residues, namely P0A7S3 (56.45%), P07017 (55.87%), P02359 (52.51%), and P46889 (52.44%). While, the proteins which exhibited a significantly low percentage of disordered residues were P24228 (5.87%) and P23872 (5.95%). Conversely, the protein sequence P23872 exhibited majorly highly flexible residues with a propensity of 64.19%. while, the highest occurrence of highly flexible residue was noted in P0AA25 (72.81%). The highest occurrence of moderately flexible residues was found in P12282 (59.03%), while the lowest was noted in P02359 (6.14%). In contrast, the lowest (zero) values in the other section were noted in sixteen moonlighting sequences (A1AGC3, A7ZUJ7, P0A7V8, P68187, P0A7E9, P0A7S3, P00805, B1LHD3, P0ADY3, Q47223, P0C0V0, P0ABS8, P02359, P0A7R5, A1AGJ5, and P0A890).

**Figure 9:**
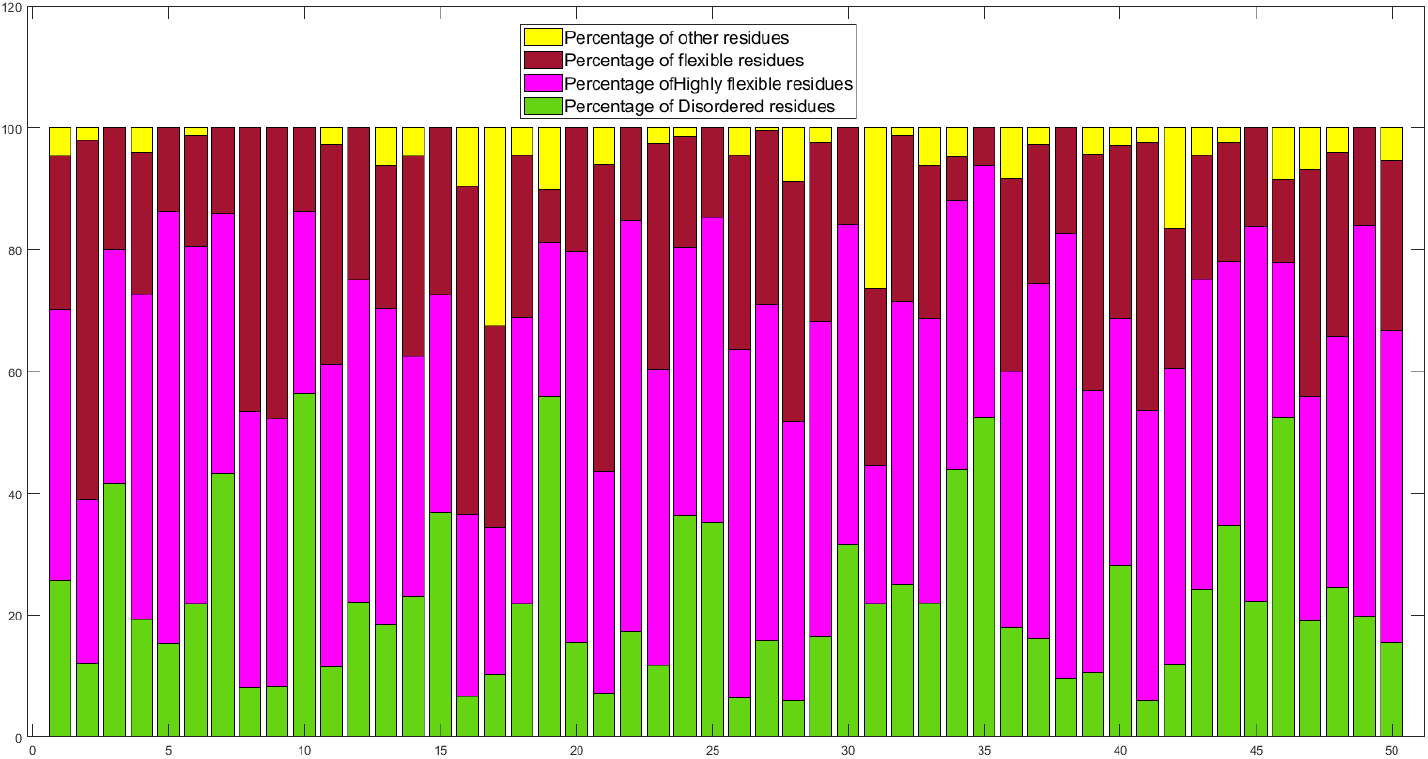
Percentages of disordered, highly flexible, moderately flexible and other residues in moonlighting proteins

#### 4.5.1. Change response sequences of disordered, highly flexible, moderately flexible, and other residues profiles

For fifty *E. coli* moonlighting proteins, we analyze the distribution of transitions between four residue types: disordered (D), highly flexible (HF), moderately flexible (MF), and other (O). Figure 10 presents the relative frequency distribution of sixteen transition types based on intrinsic disorder profiles. This analysis reveals key trends in structural flexibility, as visualized by boxplots, to distinguish between moonlighting and non-moonlighting proteins.

**Figure 10:**
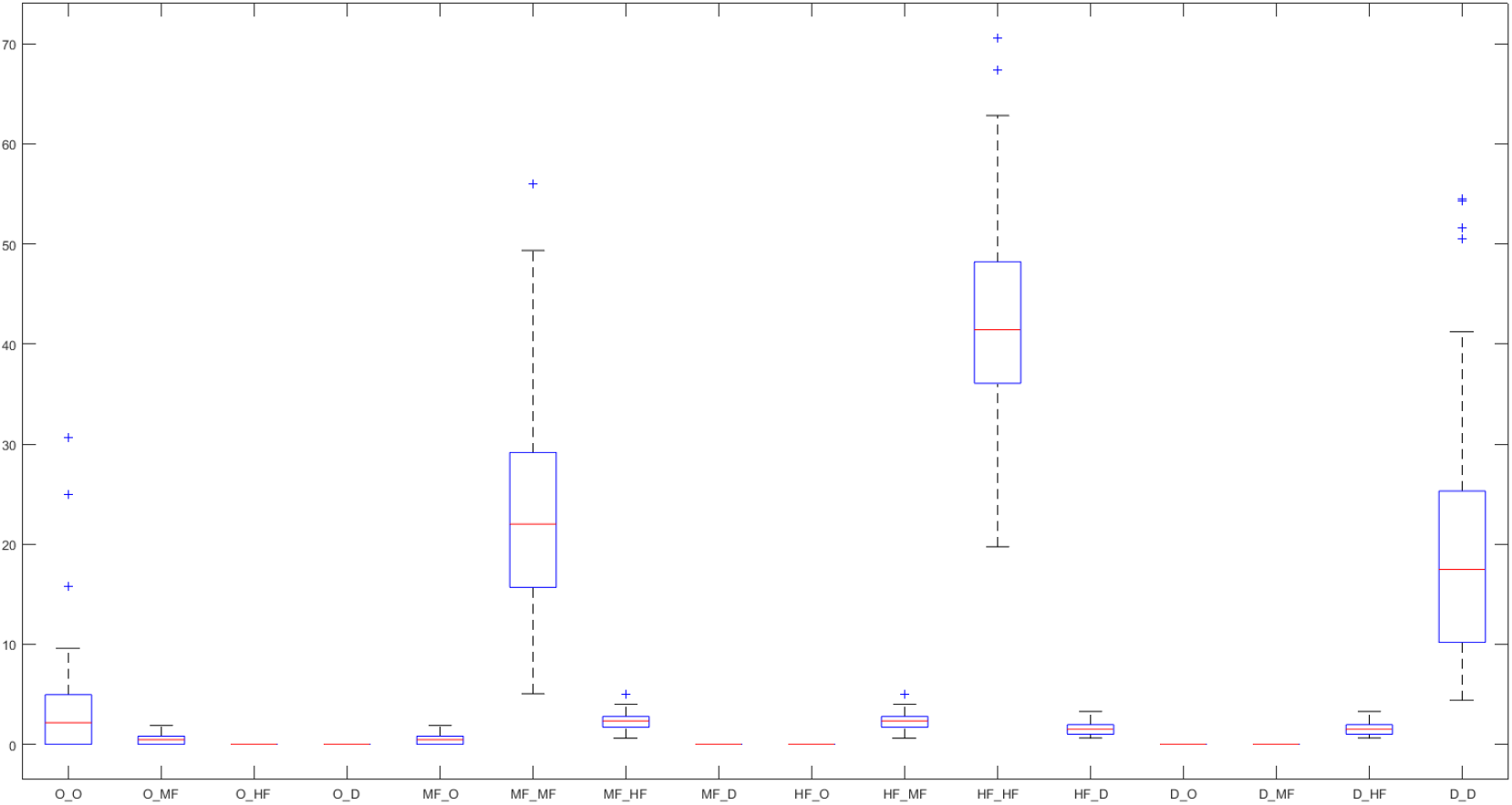
Box-plot of the percentages of disordered, highly flexible, moderately flexible and other residues changes in *E. coli* moonlighting protein sequences

P0C093 predominantly transitions between O_MF, MF_O, and MF_MF, suggesting a preference for moderately flexible states. Minimal transitions to HF or D reflect a stable structure with limited extreme flexibility. P12282 shows strong O_MF and MF_MF transitions with few shifts to HF or D, indicating a stable protein with moderate flexibility. A1AGC3 transitions between MF_O, D_MF, and D_D, suggesting dynamic flexibility with a shift between moderately flexible and disordered states. P30749 transitions mostly between MF_MF, HF_MF, and MF_D, highlighting regions of both stability and moderate flexibility. A7ZUJ7 shows minimal transitions, with a preference for MF_O and O_MF, indicating a stable, moderately flexible structure. A1AIF7 transitions from O_MF and MF_MF to D_O, with some flexibility towards disordered regions. P0A7V8 predominantly transitions between O_MF and MF_MF, with minimal shifts to HF or D. P68187 shows strong MF_MF transitions, with few changes towards HF or D, indicating a preference for moderately flexible states. P0A7E9 transitions mainly between MF_MF and MF_D, suggesting moderate flexibility with occasional shifts towards disordered states. P0A7S3 favors MF_O and O_MF transitions, with some movement towards D_MF. Minimal shifts to HF suggest a more rigid, moderately flexible structure.

None of the fifty moonlighting proteins showed transitions from disordered to moderately flexible, disordered to other, highly flexible to other, moderately flexible to disordered, other to disordered, or other to highly flexible residues. Notably, P15531 exhibited the highest self-transition (50.33%) from moderately flexible to moderately flexible, while P07355 had the highest self-transition (63.91%) from highly flexible to highly flexible. Figure 10 highlights these distinctive transition patterns among moonlighting protein sequences.

An analysis of 50 moonlighting sequences, using a distance threshold of 19, resulted in the formation of seven distinct clusters, which can be observed in the dendrogram (Figure 11 and are summarized in Table 6). The most substantial cluster, Cluster-2, consisted of 17 sequences and is highlighted by a red in Figure 11. Clusters range from small (e.g., Cluster-7 with only 2 sequences) to large (e.g., Cluster-2 with 17 sequences), suggesting that some proteins share common flexibility characteristics while others are more specialized.

**Figure 11:**
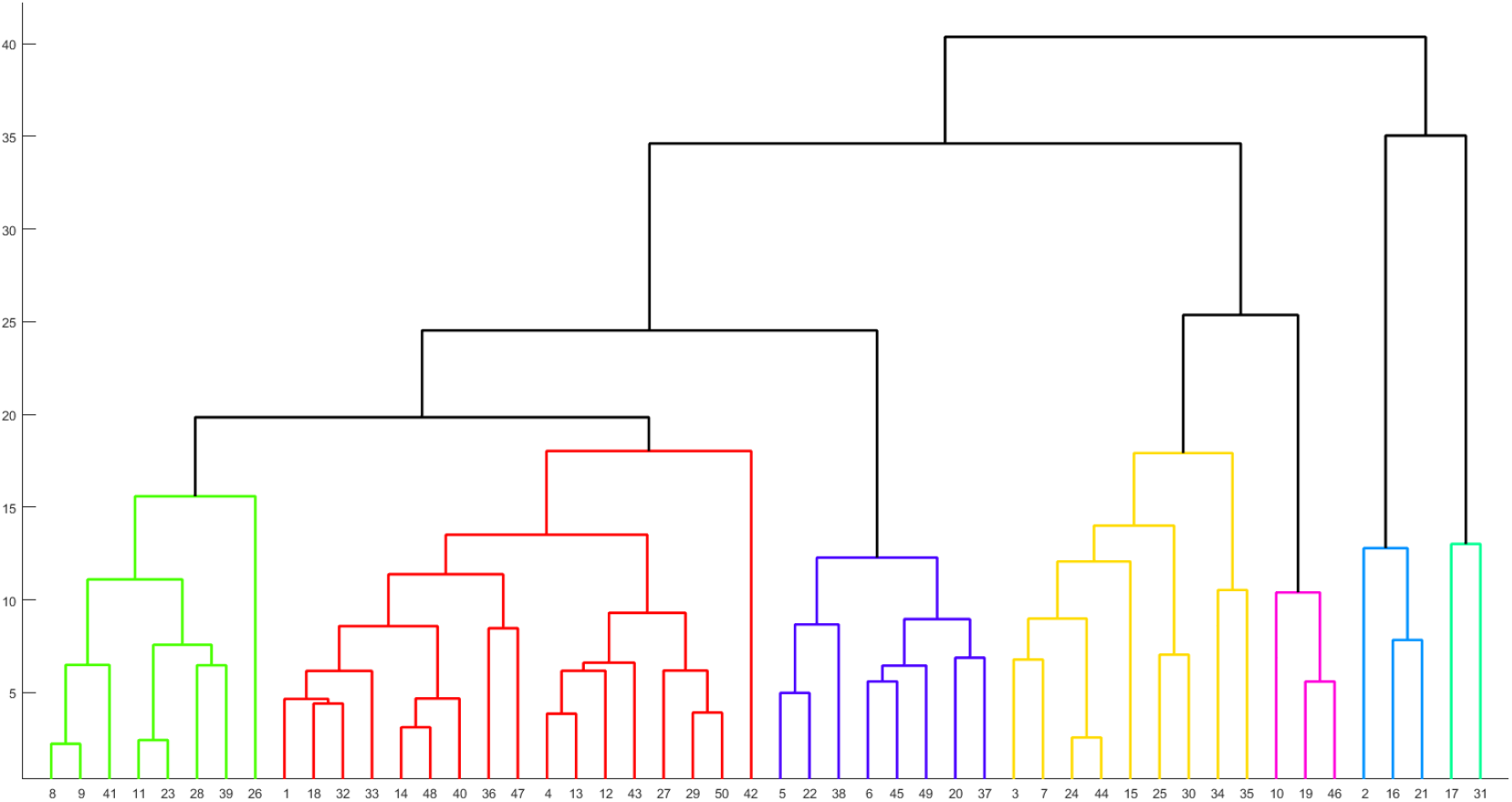
Phylogenetic relationship among the *E. coli* moonlighting proteins based on percentages of disordered, highly flexible, moderately flexible and other residues.

**Table 6:**
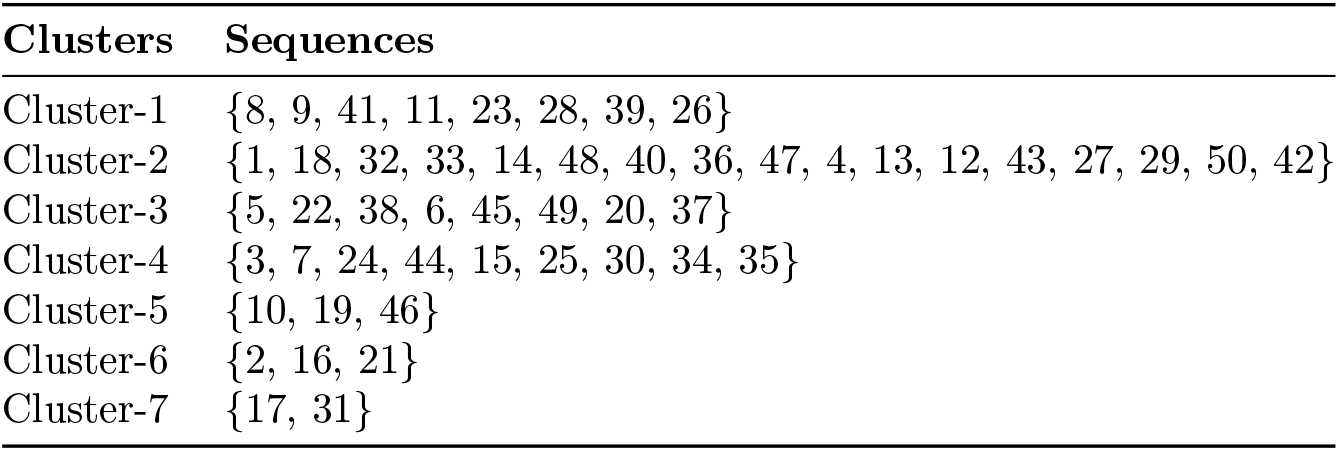
Clusters of moonlighting proteins based on the change response profiles of disordered, highly flexible, moderately flexible, and other residues in *E. coli* moonlighting proteins.

| Clusters | Sequences |
| --- | --- |
| Cluster-1 | {8, 9, 41, 11, 23, 28, 39, 26} |
| Cluster-2 | {1, 18, 32, 33, 14, 48, 40, 36, 47, 4, 13, 12, 43, 27, 29, 50, 42} |
| Cluster-3 | {5, 22, 38, 6, 45, 49, 20, 37} |
| Cluster-4 | {3, 7, 24, 44, 15, 25, 30, 34, 35} |
| Cluster-5 | {10, 19, 46} |
| Cluster-6 | {2, 16, 21} |
| Cluster-7 | {17, 31} |

### 4.6. Proximal relationships among E.coli moonlighting proteins

Seven sets of proximal sets consist of 20 *E. coli* moonlighting proteins were identified through clustering of the signature features of moonlighting protein sequences, as shown in Tables (2, 4, 5, and 6).

**Table 7:** Proximity of *E. coli* moonlighting proteins based on various quantitative signature features.

| Proximal Sets | Proximal set of *E. coli* moonlighting proteins |
| --- | --- |
| **Proximal Set 1:** | {P68187, P0A7E9, P76015} |
| **Proximal Set 2:** | {P0A9Q7, P23872} |
| **Proximal Set 3:** | {P12281, P09546, P36683, P0AF03, P27302} |
| **Proximal Set 4:** | {P68767, P31658} |
| **Proximal Set 5:** | {P0A9M0, P0A8M3, P00957, P27278} |
| **Proximal Set 6:** | {A1AGC3, P02359} |
| **Proximal Set 7:** | {P12282, P23256} |

#### 4.6.1. Typical and atypical roles of moonlighting proteins within the proximal sets

The set of proteins within each of the seven proximal groups of *E. coli* moonlighting proteins were found to be involved in a range of both common and unique functions. For each proximal group, the common and uncommon functions are outlined here.

##### Proximal Set 1

P68187 (Aldehyde dehydrogenase), P0A7E9 (Glutamate dehydrogenase), and P76015 (Acetate kinase) *Common function:* Involved in metabolic pathways (detoxification, amino acid metabolism, and energy metabolism) [43].

###### Uncommon functions

P68187: In addition to its role in detoxifying aldehydes, it is sometimes involved in cellular stress responses. It may also play a role in regulating the redox state of the cell [44].

P0A7E9: Besides its metabolic role, this enzyme has been implicated in feedback regulation in response to nitrogen availability [44].

P76015: It might also play a role in regulating acetate levels under certain environmental conditions, ensuring that metabolic pathways remain balanced [44].

##### Proximal Set 2

P0A9Q7 (Formate dehydrogenase), P23872 (Phosphoglycerate mutase)

###### Common function

Both are involved in central metabolic processes (carbon metabolism, glycolysis) [45].

###### Uncommon functions

P0A9Q7: In addition to carbon metabolism, it is important for electron transfer, as formate dehydrogenase can participate in electron transport in certain anaerobic conditions [44].

P23872: While it is essential for glycolysis, this enzyme can also be involved in gluconeogenesis, a process that occurs under stress or when the cell needs to produce glucose from non-carbohydrate sources.

##### Proximal Set 3

P12281 (Malate dehydrogenase), P09546 (Phosphoglucose isomerase), P36683 (Glutamine synthetase), P0AF03 (Ribonucleotide reductase), and P27302 (Urease)

*Common function:* Involved in metabolic biosynthesis and energy production [46].

###### Uncommon functions

P12281: While it is primarily involved in energy production (TCA cycle), malate dehydrogenase can also help in pH regulation within cells by maintaining a balance of acidic and basic molecules [47].

P09546: It can have roles in cell signaling in certain contexts, especially when glucose metabolism is linked to gene expression [44].

P36683: Beyond its role in nitrogen assimilation, glutamine synthetase has been involved in regulation of gene expression, especially genes related to nitrogen metabolism [48].

P0AF03: This enzyme doesn’t just produce the building blocks for DNA synthesis; it also contributes to DNA repair by ensuring a steady supply of deoxyribonucleotides [49].

P27302: While primarily involved in nitrogen metabolism, urease has been shown to influence virulence in pathogenic bacteria like *E. coli* by increasing ammonia production and affecting pH levels in the surrounding environment [50].

##### Proximal Set 4

P68767 (GroEL), P31658 (GroES)

###### Common function

Involved in protein folding and maintaining protein integrity [51].

###### Uncommon functions

P68767: GroEL may also have a role in stress tolerance mechanisms, as it helps to stabilize proteins under heat shock conditions [52].

P31658: Besides its chaperone role, GroES can also be involved in protein complex assembly and may influence the stability of protein-protein interactions within the cell [53].

##### Proximal Set 5

P0A9M0 (Fumarase), P0A8M3 (Succinyl-CoA synthetase), P00957 (Glyceraldehyde-3-phosphate dehydrogenase), and P27278 (Phosphoenolpyruvate carboxykinase)

###### Common function

Involved in central metabolic pathways (citric acid cycle, glycolysis, gluconeogenesis) [54].

###### Uncommon functions

P0A9M0: Apart from its metabolic role, fumarase can also be involved in cellular oxidative stress response as it helps to regulate the redox state of cells [55].

P0A8M3: This protein plays a role in bacterial adaptation to different environmental conditions by regulating metabolic fluxes [56].

P00957: GAPDH is well known for its glycolytic function, but it is also involved in nuclear processes such as DNA repair and may function as a moonlighting protein in these pathways [57].

P27278: In addition to its role in gluconeogenesis, this protein is involved in acid-base regulation during high metabolic activity [44].

##### Proximal Set 6

A1AGC3 (Cysteinyl-tRNA synthetase), P02359 (RNA polymerase sigma-70 factor)

###### Common function

Involved in protein synthesis and gene expression regulation [58].

###### Uncommon functions

A1AGC3: Beyond its tRNA synthetase function, it can participate in the regulation of translation efficiency during stress conditions [**?**].

P02359: Sigma-70 is crucial for transcription, but it can also have roles in stress response regulation by altering the transcriptional profile of the cell during environmental changes [59].

##### Proximal Set 7

P12282 (Glycogen synthase), P23256 (Glucokinase)

###### Common function

Involved in carbohydrate metabolism and energy storage [60].

###### Uncommon functions

P12282: Apart from its role in glycogen biosynthesis, it is involved in regulation of cellular energy balance, especially under fluctuating glucose levels [61].

P23256: Beyond glucose phosphorylation, glucokinase plays a role in glucose sensing and regulation of insulin secretion in higher organisms. In *E. coli*, it may contribute to the regulation of glycolytic flux [62].

The seven proximal sets of *E. coli* moonlighting proteins perform a variety of common and uncommon functions. Common roles include involvement in central metabolic processes like energy production, amino acid, and carbohydrate metabolism (e.g., aldehyde dehydrogenase, phosphoglycerate mutase, and malate dehydrogenase). Several proteins are crucial for protein synthesis (e.g., cysteinyl-tRNA synthetase, RNA polymerase sigma-70) and protein folding (e.g., GroEL, GroES). Uncommon functions include participation in stress responses, oxidative stress regulation, and acid-base balance. Some proteins, such as glutamine synthetase and glyceraldehyde-3-phosphate dehydrogenase, also aid in gene regulation and DNA repair [63]. Additionally, glucokinase and glycogen synthase help regulate energy balance and glucose sensing, showcasing the diverse moonlighting roles these proteins play in bacterial cells [64].

## 5. Discussion

This study provides a quantitative analysis of fifty *E. coli* moonlighting proteins across multiple dimensions, leading to the identification of distinct clusters based on the extracted features, as detailed in Section 4. The analysis of amino acid distribution revealed substantial variability in the composition and frequency of amino acids across the sequences. Alanine was predominant, with the highest median frequency and standard deviation, indicating its significant role in these proteins. Although Alanine is a key building block in bacterial cells, excessive accumulation can be toxic. *E. coli* moonlighting proteins, which have a higher proportion of Alanine, are regulated by the Alanine exporter, acting as a safety valve to maintain safe intracellular levels and prevent toxicity [65, 66]. Conversely, Cysteine had the lowest median and standard deviation, suggesting a more specialized function. The absence of certain amino acids, such as Tryptophan and Cysteine, further emphasizes the variability in functional requirements [67]. The clustering analysis imply sequence Q47223 has a unique relative frequency of amino acids, differentiating it from remaining proteins. In case of human moonlighting proteins, the highest frequency of Leucine was observed in most number of sequences, while the highest frequency of Alanine was observed in most number of sequences in *E. coli* [22].

The longest poly-strings of length 6 were found only in two sequences, suggesting that while long repeats are possible, they are rare. One of the key differences that was noted between human and *E.coli* in poly-string frequency is presence of length 5 with maximum length 14 in human moonlighting proteins, while absence of length 5 with maximum length 6 in *E.coli* [22]. These patterns suggest that repetitive sequences are selectively incorporated into moonlighting proteins for specific functions, with the length and type of repeats influenced by structural and functional requirements [68].

The polar and non-polar residue analysis revealed a higher proportion of non-polar residues, with an average polar-to-non-polar ratio of 0.89 indicating a preference for non-polar interactions, which may enhance protein stability and functional versatility. Notably, P69786 exhibited the lowest ratio of 0.5 and consequently highest percentage of non-polar to non-polar changes, stand out in terms of biochemical properties. The average ratio of polar to non-polar residues was 1.0061 *±* 0.2462 and the highest ratios of polar-to-non-polar residues was 2.384 in human moonlighting proteins while the highest ratio was 1.28 for *E.coli* [22].

The analysis of acidic, basic, and neutral residue profiles revealed variability in their distribution may reflect the structural constraints or functional specificity required for certain proteins to perform their moonlighting functions effectively. P12282 did not have any basic to basic and acidic to basic changes. Q47223 did not reveal any basic to acidic, acidic to acidic and basic to basic. P0ABS8 contained neither acidic to acidic nor basic to basic changes. Percentages of neutral residues spanned from 50.2 to 93.2 in human moonlighting proteins while the same varied from 67.47 to 88.04 in *E.coli* [22]. The highest percentage of basic residue was 29.41 and 22.58 in human and *E.coli* moonlighting proteins respectively [22]. NN changes varied from 23.25% to 88.95% and 42.92% to 76.5% in human and *E.coli* moonlighting proteins respectively [22].

The intrinsic disorder analysis of *E. coli* moonlighting proteins revealed significant variability in the proportion of disordered, highly flexible, and moderately flexible residues. Notably, these flexibility profiles, combined with distinct change responses, suggest that different moonlighting proteins exhibit varying degrees of structural flexibility, likely influencing their functional adaptability [69, 70]. It was noticed that there was a significant difference between the highest percentage of disordered residues in human (100%) and *E.coli* (56.45%) moonlighting proteins, as well as average percentage of the same of former was significantly more than that of latter indicating difference between structural stability [22]. The clustering highlights the structural diversity among *E. coli* moonlighting proteins, further supporting the idea that flexibility and disorder play crucial roles in their functional versatility.

Finally, the cumulative clustering of the *E. coli* moonlighting proteins revealed seven distinct proximal sets. Proximal sequences contribute to central metabolic processes such as glycolysis and energy production, while also play roles in stress responses, oxidative regulation, and gene expression. These findings underscore the versatility of *E. coli* moonlighting proteins, demonstrating their capacity to adapt to varying environmental conditions and contribute to a wide range of cellular functions [71]. The diversity in structural and functional characteristics of these proteins highlights their importance in enabling bacterial adaptation, stress resilience, and metabolic homeostasis [72].

## Acknowledgments

The authors sincerely thank Mr. Ashim Dhar and Mrs. Piyali Maitra for their logistical support in the laboratory. A special thanks is extended to Mr. Arindam Samanta for his valuable and insightful comments. The authors also express their gratitude to the Indian Statistical Institute (ISI/TAC/PROJECT-1/2024-25) for their financial support. Notably, feature extractions and subsequent computations were carried out in MATLAB (2024b).

## Author contributions statement

DN, SSH, AG, and VNU formulated the problem and designed the theoretical experiments. DN, SSH, MS, VNU, and Ankita Ghosh carried out the experiments and performed the analyses. DN, SSH, VNU, MS, and Ankita Ghosh drafted the initial manuscript. All authors contributed to reviewing and editing the manuscript. SSH and AG supervised the overall project. All authors reviewed, checked, and approved the final manuscript.

